# A maize lipid droplet-associated protein is modulated by a virus to promote viral multiplication and infection

**DOI:** 10.1101/2023.02.02.526882

**Authors:** Siyuan Wang, Siqi Li, Xinyu Wang, Xi Sun, Mingshuo Xue, Dianping Di, Aihong Zhang, Yongjiang Zhang, Yiji Xia, Tao Zhou, Zaifeng Fan

## Abstract

Pathogen infection induces massive reprogramming of host primary metabolism. Lipid and fatty acid metabolism is generally disrupted by pathogens and co-opted for their proliferation. Lipid droplets (LDs) that play important roles in regulating cellular lipid metabolism are utilized by a variety of pathogens in mammalian cells. However, the function of LDs during pathogenic infection in plants remains unknown. We show here that infection by rice black streaked dwarf virus (RBSDV) affects the lipid metabolism of maize, which causes elevated accumulation of C18 polyunsaturated fatty acids (PUFAs) leading to viral proliferation and symptom development. Overexpression of one of the two novel LD-associated proteins (LDAPs) of maize (ZmLDAP1 and ZmLDAP2) induces LD clustering. The core capsid protein P8 of RBSDV interacts with ZmLDAP2 and prevents its degradation through the ubiquitin-proteasome system mediated by a UBX domain-containing protein, PUX10. In addition, silencing of the *ZmLDAP2* down-regulates expression of fatty acid desaturase genes in maize, leading to a decrease in C18 PUFAs levels and suppression of RBSDV accumulation. Our findings reveal that the plant virus recruits LDAP to regulate cellular fatty acid metabolism to promote viral multiplication and infection. These results expand the knowledge of the LD functions and viral infection mechanism in plants.

**One Sentence Summary:** Rice black streaked dwarf virus recruits a lipid droplet-associated protein to regulate cellular fatty acid metabolism for promoting viral multiplication and infection.

## Introduction

Plants exposed to external stimuli constantly show intricate changes of primary metabolism, involved in photosynthesis and metabolisms of carbohydrates, amino acids and lipids. These metabolic processes are utilized by plant pathogens for their proliferation (Bolton, 2009; Rojas et al., 2014). During virus infection, cellular lipids provide energy for metabolism, participate in multiple defense signal cascades, and serve as structural components of intracellular membranes for replication of some viruses (Heaton and Randall, 2011; Nagy and Pogany, 2011; Wang, 2004). Plant viruses interfere with lipid and fatty acid (FA) metabolism to promote changes in membrane fluidity and plasticity which are required for the proper assembly of viral replication complexes (Heaton and Randall, 2011; Llave, 2016).

Linoleic acid (LA, C18:2) and α-linolenic acid (ALA, C18:3) are common polyunsaturated FAs (PUFAs) in plant lipids, and regarded as important regulators in plant defense (Lim et al., 2017). For example, increased levels of LA and ALA result in higher resistance to *Pseudomonas syringae* in tomato (Yaeno et al., 2004). Elevated levels of ALA have been reported in tobacco infected with tobacco mosaic virus (Choi et al., 2006). Accumulation of tobacco rattle virus was reduced in *fad2 Arabidopsis thaliana* mutants that contained reduced levels of PUFAs (LA and ALA) (Fernández-Calvino et al., 2014). These studies suggest that C18 PUFAs play important roles in infection of plants by various pathogens.

Lipid droplets (LDs) are universal organelles used for neutral lipids storage in eukaryotic cells. Besides the basic functions in energy storage, LDs also play roles in modulation of FAs trafficking, lipid signaling and host defense against intracellular pathogens such as viruses (Welte, 2015). The surface of LDs is the most inert structure among all physical or membranous structures inside a cell and thus suitable for use as a refuge for proteins (Huang, 2018). As a result, LD surface can be taken advantage of by some mammalian viruses, such as hepatitis C virus (HCV), dengue virus and rotavirus, which assemble virus particles on LDs with minimal disturbance to the host cells (Barba et al., 1997; Cheung et al., 2010; Randall, 2018; Roingeard and Melo, 2017). For example, rotavirus major viroplasm proteins, NSP2 and NSP5, colocalize with perilipin A, an LD structural protein, and form complexes with cellular LDs in early infection (Cheung et al., 2010). Plant LDs occur not only in storage organs (such as seeds, pollens, flower tapetum and fruit mesocarp) serving as storage warehouses, but also in non-storage vegetative organs (such as leaves, stems and roots) serving as detoxification refuges (Huang, 2018). LDs in leaves are induced when plants are subjected to abiotic or biotic stresses, during senescence, at the end of the dark period in a diurnal cycle, or exposed to a pathogenic fungus (Gidda et al., 2016; Shimada and Hara-Nishimura, 2015; Shimada et al., 2014). However, it has not been reported if plant LDs play any role in viral infection.

LDs are decorated by numerous specific proteins, which are varied among cell and tissue types. LD proteins can be classified into two groups based on how they are targeted to LDs (Kory et al., 2016). In plants, oleosins are the most prominent structural LD proteins in storage tissues but are absent in vegetative tissues (Kim et al., 2002). Several proteins with enzymatic activities, like caleosin, dioxygenase and steroleosin, are also found to be targeted to plant LDs (Shimada et al., 2014). In recent years, LD-associated proteins (LDAPs) have been identified in non-seed tissues such as leaf and fruit tissues (Gidda et al., 2013; Horn et al., 2013; Kim et al., 2016). Horn et al. identified a class of LDAPs in various plants using phylogenetic analysis (Horn et al., 2013). This kind of LD proteins is thought to be the main structural proteins of leaf LDs and participate in cellular stress defenses (Huang, 2018). *Arabidopsis* LDAPs were shown to play positive roles in LD biogenesis, tissue growth, cell proliferation, cell wall organization, and drought stress responses (Kim et al., 2016). However, the functions of plant LDAPs in viral infection remain elusive.

Plant UBX (ubiquitin regulatory X) domain-containing protein10 (PUX10), which is a member of the plant UBX domain-containing (PUX) protein family, is localized to LDs via its hydrophobic domain (Kretzschmar et al., 2018). Studies in *Arabidopsis* seeds and tobacco pollen tubes demonstrated that PUX10 could recruit the AAA (ATPase associated with various cellular activities) ATPase CDC48 (cell division cycle 48) to ubiquitinated LD proteins (oleosins) by interacting with its UBX and UBA (ubiquitin-associated domain) domains, respectively, thus facilitating oleosins segregated from LDs into degradation pathway (Deruyffelaere et al., 2018; Kretzschmar et al., 2018). Besides, the PUX10 mutation increased the ratio of C18:3 to C18:2 FAs in *Arabidopsis* seeds, suggesting that PUX10 plays a role in PUFA metabolism (Deruyffelaere et al., 2018).

Rice black-streaked dwarf virus (RBSDV) belongs to the genus *Fijivirus* in the family of *Reoviridae*. RBSDV often results in severe losses in the production of maize (*Zea mays* L.), rice, wheat, and other graminaceous crops (Bai et al., 2001; Wu et al., 2020). The infection in maize plants causes rough dwarf disease, exhibiting severe growth abnormalities, including plant dwarfing, dark green leaves, and waxy white tumors on sheaths and veins of the abaxial surface of the leaves (Favali et al., 1980; Shen et al., 2016). The genomes of RBSDV consist of ten linear double-stranded RNA (dsRNA) segments (S1-S10), each encoding one or two (for S5, S7, and S9) proteins (Wang et al., 2003; Zhang et al., 2001). Among the 13 viral proteins, P8 and P10 are the core- and outer-capsid protein, respectively (Liu et al., 2007a, b). P8 is reported to localize to the nucleus of insect vector and plant cells and repress transcription in tobacco suspension cells (Liu et al., 2007a). Our previous study revealed that P8 interacted with maize AKINβγ, the regulatory subunit of the SNF1-related protein kinase 1 (SnRK1) complex, thus regulating primary carbohydrate metabolism of host plant in response to RBSDV infection (Li et al., 2020). In addition, SP8 encoded by southern rice black-streaked dwarf virus, a closely related fijivirus, was reported to interfere with the dimerization of auxin response factor 17 of rice thus disturbing the auxin signaling pathway and facilitating virus infection (Zhang et al., 2020). It has been reported that the P5-1, P6 and P9-1 proteins constitute viroplasm, where viral replication and assembly occur (Akita et al., 2012; Sun et al., 2013; Wang et al., 2011). Recent reports showed that P5-1 hinders the ubiquitination activity of SCF E3 ligases leading to inhibiting jasmonate (JA) signaling to benefit viral infection in rice (He et al., 2020). However, the host primary metabolic pathways disturbed by RBSDV infection remain to be elucidated.

Here, we describe the use of metabolomic approaches to gain new insights into the metabolic changes that occur in maize in response to RBSDV infection. The results showed that two C18 PUFAs, ALA and LA, were significantly induced upon RBSDV infection. Further analyses showed that exogenous ALA treatment significantly enhanced maize susceptibility to RBSDV infection. In this research, we have identified two LDAPs, ZmLDAP1 and ZmLDAP2, and shown that the ZmLDAP2 transcript level is up-regulated by RBSDV infection. We found that RBSDV P8 interacts with ZmLDAP2 and associates with the clustered LDs induced by ZmLDAP2 overexpression. Knockdown of *ZmLDAP2* in maize inhibits synthesis of LA and ALA and reduces susceptibility of maize to RBSDV infection. We also found that ZmLDAP2 is targeted by ZmPUX10 for degradation through ubiquitin-proteasome system, and the degradation is specifically inhibited by P8. Together, our findings demonstrate that RBSDV interferes with the degradation of the maize LD protein (so that it can utilize LDs) to regulate the C18 PUFAs metabolism for enhancing viral multiplication and infection.

## Results

### RBSDV infection alters the metabolite composition in maize leaves

To gain insights into the metabolic response of maize to RBSDV infection, we performed a large-scale metabolomic analysis of maize seedlings at 10 and 28 days post-inoculation (dpi), which represent the pre-symptomatic phase (no symptom but viruses were detected by RT-PCR) and the symptomatic phase (typical symptoms of RBSDV including the stunted phenotype, dark green leaves, and waxy white tumors on leaves), respectively (Fig. 1A). Six samples of both RBSDV-infected and mock-inoculated plants at each phase were analyzed by gas chromatography coupled with a Pegasus HT time-of-flight mass spectrometer (GC-TOF-MS). This analysis identified a total of 279 primary metabolites (Supplemental Table S1), and metabolite composition was subsequently investigated and compared between RBSDV infection groups and mock groups. First, an unsupervised clustering method, principal component analysis (PCA), and a supervised clustering method, orthogonal projections to latent structures-discriminant analysis (OPLS-DA), were performed on the GC-MS data, showing clear separation between RBSDV-infected and mock-inoculated groups (Supplemental Fig. S1). Univariate statistical analysis (Student’s *t*-test) was performed to detect metabolites significantly affected by RBSDV infection. The responsive metabolites were isolated on the basis of VIP > 1 and *P*-value < 0.05.

**Figure 1.**
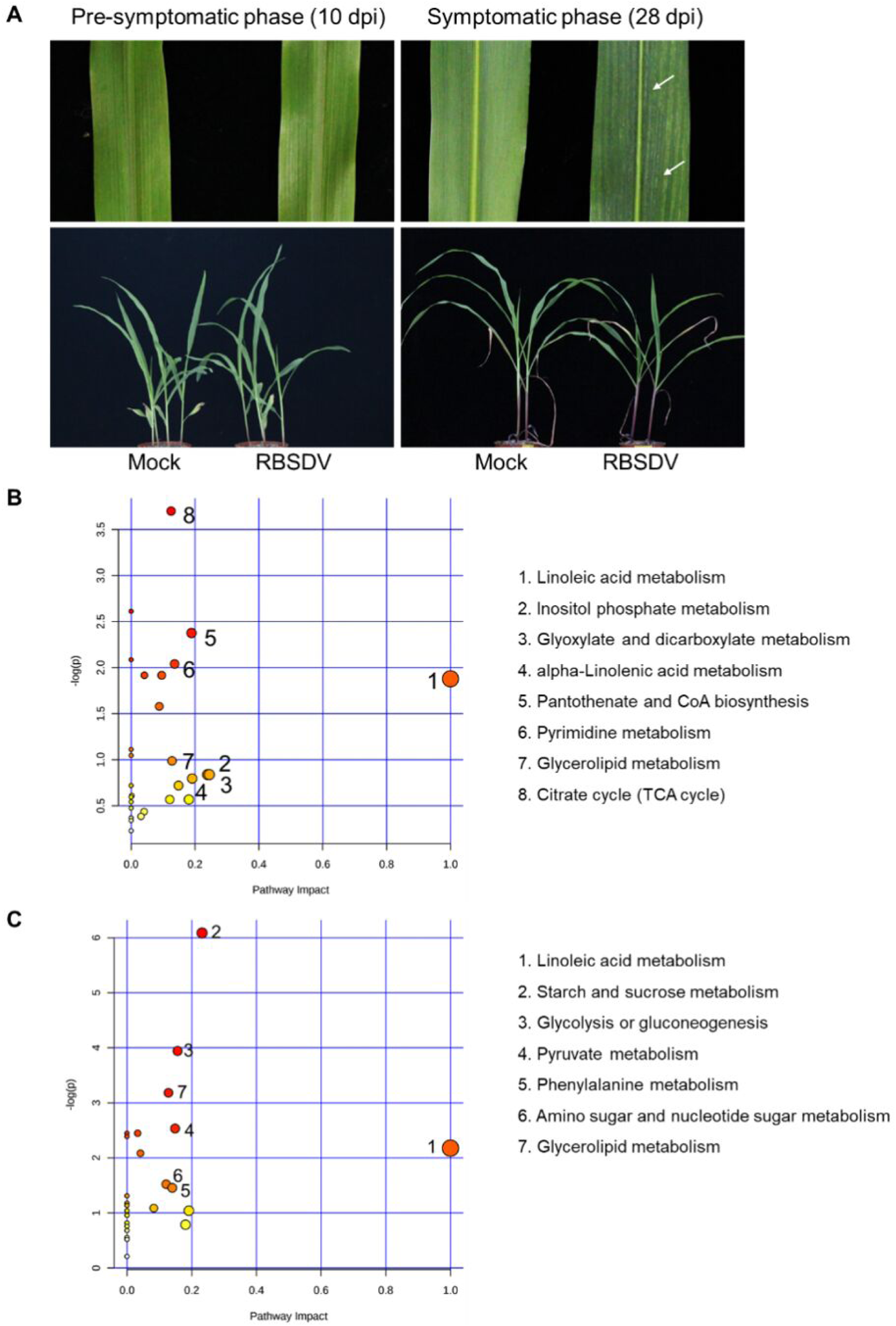
Differentially regulated metabolites in maize leaves infected with RBSDV. **A,** Infection phases and the symptoms on RBSDV- and mock-inoculated maize seedlings at 10 dpi (pre-symptomatic phase) and 28 dpi (symptomatic phase). The white arrows indicate the waxy white streaks on leaves. **B and C**, Pathway enrichment analysis performed using the significantly different metabolites between RBSDV-infected and mock-inoculated groups at 10 dpi (B) and 28 dpi (C). Bubble color represents the *P* value: deeper colors represent smaller *P* values, indicating larger differences. The size of the bubble represents the impact of the pathway during topological analysis. A larger size represents higher impact.

Comparing the RBSDV-infected groups and mock-inoculated groups in the two infection phases separately, we found that there were 53 metabolites differentially expressed at the pre-symptomatic phase (10 dpi) and 41 metabolites at the symptomatic phase (28 dpi) (Supplemental Table S2 and S3). The detailed differentially regulated metabolites are listed in a visualized square (Supplemental Fig. S2). There are eight differentially expressed metabolites that are present in both phases. Furthermore, the pathway analysis was performed by MetaboAnalyst (http://www.metaboanalyst.ca) based on the KEGG pathway database (Fig. 1B, C). From the enrichment analysis results (as shown in the Y-axis), the citrate cycle is the most significantly affected pathway at the pre-symptomatic phase, and the starch and sucrose metabolism is the most significantly affected pathway at the symptomatic phase. Pathway impact results (as shown in the X-axis) indicated that the linoleic acid metabolism presented the highest impact at the both two phases. Together, we postulated that the linoleic acid metabolism pathway is markedly perturbed by RBSDV infection.

### ALA plays positive roles during RBSDV infection in maize

According to the GC-MS data, two C18 PUFAs, ALA and LA, showed a significant increase in response to RBSDV infection. The levels of ALA were up-regulated 3.4-fold in the pre-symptomatic phase and 3.1-fold in the symptomatic phase comparing with mock-inoculated control (Fig. 2A). The levels of LA had a 2.7-fold increase in the pre-symptomatic phase and 1.5-fold in the symptomatic phase (Fig. 2B).

**Figure 2.**
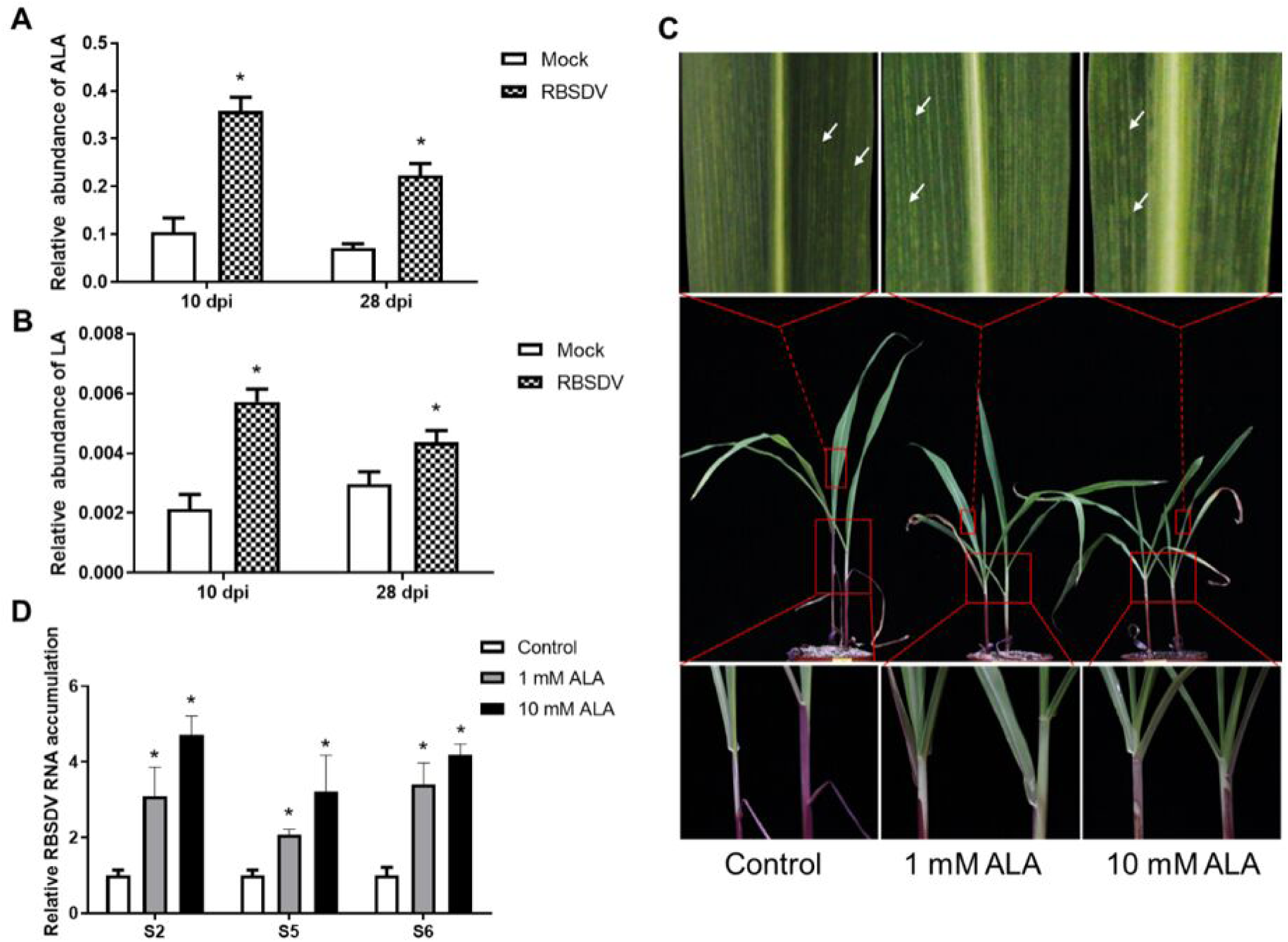
α-Linolenic acid (ALA) plays a positive role in RBSDV infection. **A and B**, Effect of RBSDV infection on ALA (A) and linoleic acid (LA) (B) levels in maize. Contents of ALA and LA were quantified by GC-TOF-MS on RBSDV-infected and mock-inoculated leaves. Data were analyzed using two-tailed Student’s *t*-test and bars represent standard error of the mean (SEM) (n=6). The asterisks represent statistical significance (*, *P* < 0.05). **C,** RBSDV symptoms affected by exogenous ALA treatment on RBSDV-infected maize. ALA or DMSO (as the control) were applied 24 h after virus inoculation. The phenotypes were observed at 17 dpi. The white arrows indicate the waxy white streaks on leaves. **D,** RT-qPCR results showing the relative expression of viral RNA (three different RBSDV genomic RNA segments S2, S5, and S6) in ALA-applied and control maize leaves at 17 dpi. At least 30 seedlings were used for each treatment, and three experiments were repeated. Error bars represent SEM calculated from at least six plants (*, *P* < 0.05, two-tailed Student’s *t*-test).

To investigate the detailed association between the C18 PUFAs and RBSDV infection, the expression levels of ALA and LA biosynthetic genes were analyzed by reverse transcription-coupled quantitative PCR (RT-qPCR) in mock- or RBSDV-inoculated plants. Omega-6 fatty acid desaturase ZmFAD2a, ZmFAD2b (cytoplasm-localized) and ZmFAD6 (chloroplast-localized) are reported to play roles in LA biosynthesis; omega-3 fatty acid desaturase ZmFAD3a, ZmFAD3b (cytoplasm-localized), ZmFAD7 and ZmFAD8 (chloroplast-localized) participate in ALA biosynthesis (Yin et al., 2018). The transcript accumulation of omega-6 fatty acid desaturase gene *ZmFAD2a* was significantly increased to two-fold in RBSDV-infected than mock-inoculated plants during the both phases. The transcript accumulation of omega-3 fatty acid desaturase genes *ZmFAD3a* and *ZmFAD3b* were significantly up-regulated by RBSDV to four-fold at the pre-symptomatic phase, while *ZmFAD7* and *ZmFAD8* were down-regulated by RBSDV infection (Supplemental Fig. S3). These results showed that the cytoplasmic C18 FA desaturation pathway was significantly activated by RBSDV infection, which might explain the accumulation of ALA and LA in response to viral infection. As the level of ALA was higher than LA in RBSDV-infected plants, ALA was selected for further study.

To study the impact of ALA on RBSDV infection, ALA was applied exogenously to RBSDV-infected seedlings. Maize seedlings were treated with ALA (1 mM or 10 mM) or DMSO as control 24 hours after inoculated with RBSDV for three days. When the waxy white streaks symptom began to appear on the control maize leaves (at 17 dpi), we observed the phenotype and analyzed the accumulation of viral RNAs (three different RBSDV genomic RNA segments S2, S5 and S6) by RT-qPCR (Fig. 2C, D). ALA-treated plants exhibited more stunted growth, shortened internode, as well as significantly higher levels of viral RNA than that in the control plants. These results suggested that ALA promotes RBSDV infection in maize seedlings.

### Genes encoding LD-associated proteins are differentially expressed in maize in response to RBSDV infection

The cellular organelle LDs are essential for lipid metabolism and especially for the level of free FAs (Olzmann and Carvalho, 2019). The research on identifying LD proteins in non-seed plant tissues predicted potential genes encoding LDAPs in several plants by phylogenetic analysis (Horn et al., 2013). We named the two genes encoding candidate LDAPs of maize as *ZmLDAP1* and *ZmLDAP2* (on the chromosomes 7 and 8 of maize), respectively.

In order to determine the expression profiles of the two genes in maize, the transcription levels of *ZmLDAP1* and *ZmLDAP2* were assessed by RT-qPCR in different tissues (leaf, leaf sheath and root) at one, two and three weeks after planting, respectively. The results demonstrated that although *ZmLDAP1* and *ZmLDAP2* are expressed throughout the seedling stage, they are differentially expressed in different tissues. *ZmLDAP1* showed the lowest expression in root, and *ZmLDAP2* was least expressed in leaf sheath (Fig. 3A, B).

**Figure 3.**
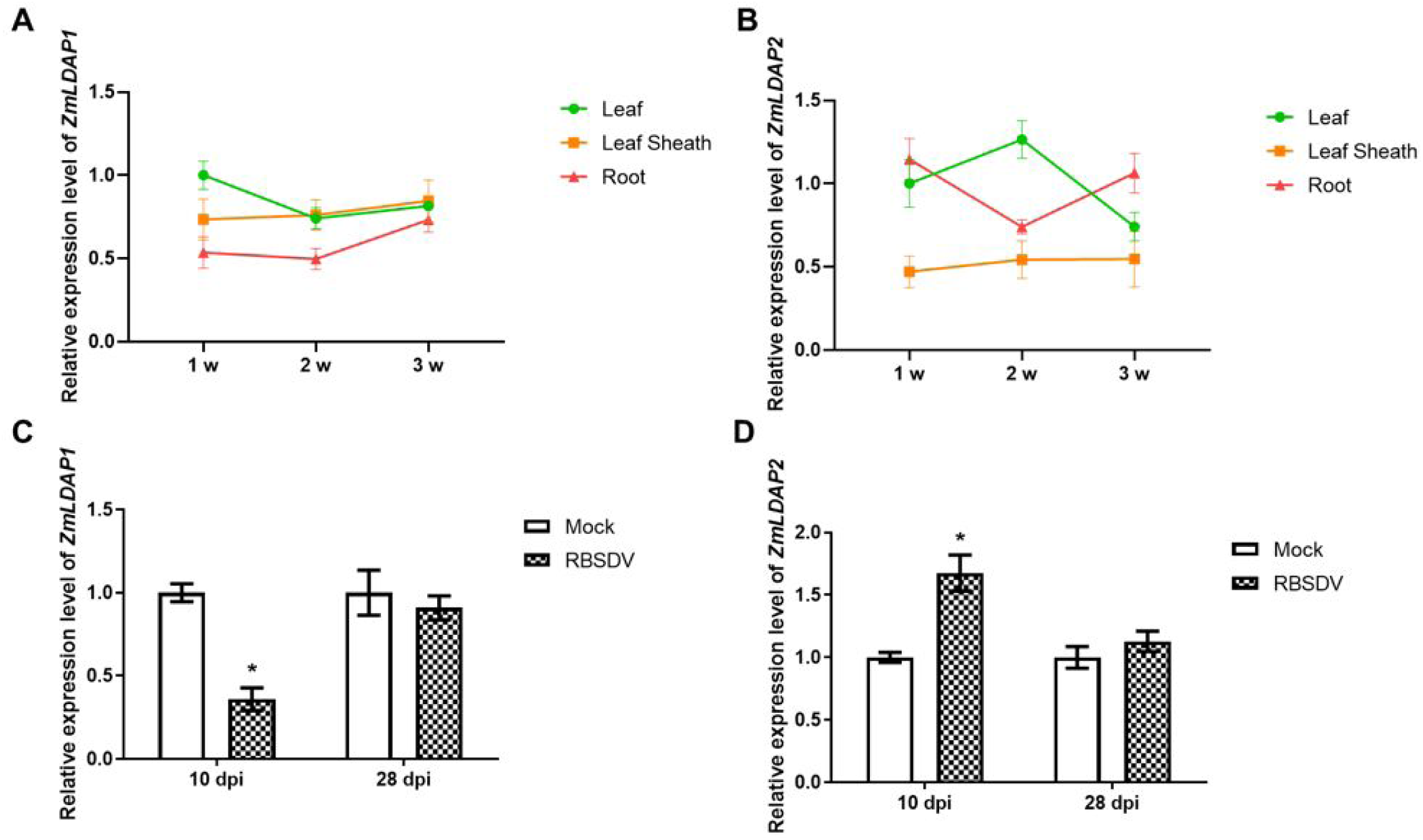
*ZmLDAP*s exhibit differential expression levels in different maize tissues and in response to RBSDV infection. **A and B**, The expression of *ZmLDAP1* (A) and *ZmLDAP2* (B) in different tissues of maize at 1 week, 2 weeks, and 3 weeks post planting. The transcript levels of *ZmLDAP1* and *ZmLDAP2* were determined through RT-qPCR using total RNA extracted from leaves (the upper newly emerged leaves), leaf sheaths and roots. Relative amounts were calculated with respect to the expression level in leaves at 1 week that were arbitrarily set at 1. Error bars represent SEM calculated from at least three plants. **C and D**, RT-qPCR results showing the relative expression levels of *ZmLDAP1* (C) and *ZmLDAP2* (D) in RBSDV-infected and mock-inoculated maize leaves (the upper newly emerged leaves) at both infection phases. Experiments were repeated three times. Error bars represent SEM calculated from at least three plants (*, *P* < 0.05, two-tailed Student’s *t*-test).

Furthermore, *ZmLDAP*s showed different expression levels during RBSDV infection. The relative transcription level of *ZmLDAP1* was significantly decreased to *ca.* 0.4-fold at 10 dpi, while *ZmLDAP2* was significantly increased to *ca.* 1.6-fold at 10 dpi (Fig. 3C, D), suggesting the different roles of ZmLDAPs in response to RBSDV early infection. The significant changes of *ZmLDAP*s at the pre-symptomatic phase was corresponding to the accumulation of C18 PUFAs and cytoplasmic *ZmFAD*s induced by RBSDV.

### RBSDV core capsid protein P8 interacts with ZmLDAP2

To determine whether the ZmLDAP1 and ZmLDAP2 proteins could associate with LDs in plant cells, the proteins tagged with GFP were transiently expressed in maize protoplasts and observed by confocal microscopy. GFP-ZmLDAP1 and GFP-ZmLDAP2 were found co-localized with Nile Red-stained LDs in large irregular structures (Fig. 4A; Supplemental Fig. S4). However, the LDs were separated in the cytoplasm in non-transfected protoplasts (Supplemental Fig. S4). This suggests that overexpression of ZmLDAP1 or ZmLDAP2 protein could promote LD aggregation.

**Figure 4.**
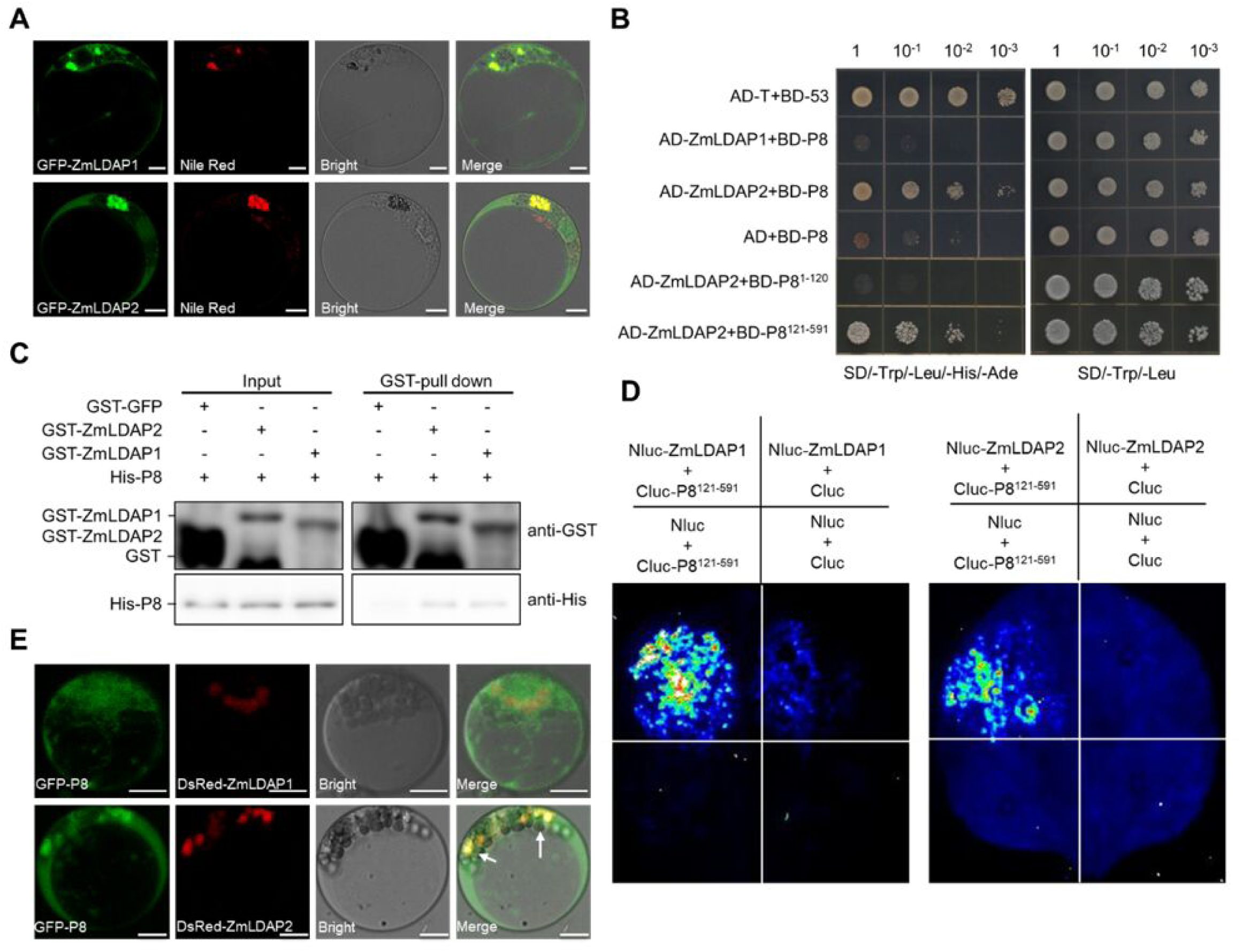
ZmLDAP2 interacts with RBSDV P8. **A,** ZmLDAP1 and ZmLDAP2 localize to LDs and induce LD clustering in maize protoplasts. GFP-ZmLDAP1 or GFP-ZmLDAP2 was transiently expressed alone in maize protoplasts. LDs were stained with Nile Red. Scale bars, 10 μm. **B,** The Y2H assay illustrates the interaction between RBSDV P8 and ZmLDAP proteins. P8^1-120^ contains a predicted nuclear localization signal domain, and P8^121-591^ contains a predicted P-loop NTPase. The different combinations of constructs transformed into yeast cells were grown on selective media SD/-Trp/-Leu, and interactions were tested with SD/-Trp/-Leu/-His/-Ade media. Pictures were taken after incubation at 30°C for 3 days. **C,** *In vitro* GST-pull-down assays demonstrating the interaction of P8 with ZmLDAP1 and ZmLDAP2. His-P8 protein was pulled down with GST-ZmLDAP1 and GST-ZmLDAP2, and further detected with anti-His antibodies. **D**, Interactions between P8^121-591^ and ZmLDAP1 or ZmLDAP2 detected by LCI assays in *N. benthamiana*. The *Agrobacterium* cells carrying the indicated constructs was infiltrated into *N. benthamiana* leaves which were subjected to fluorescence imaging at 3 days post agroinfiltration (dpai). **E,** The subcellular co-localization of GFP-P8 and DsRed-ZmLDAP1 or DsRed-ZmLDAP2 in maize protoplasts. The white arrows indicate merged fluorescence, which was produced from the overlapping of two proteins. Scale bars, 10 μm.

Previous reports indicated that LDs play important roles in viral protein assembly and virus replication in animal cells (Barba et al., 1997; Cheung et al., 2010; Laufman et al., 2019; Randall, 2018). The studies in the mammalian rotavirus, a member of the *Reoviridae* family, demonstrated that LDs associate with viroplasm where virus replication occurs (Cheung et al., 2010; Criglar et al., 2020). Therefore, we assessed whether RBSDV replication was associated with LDs in plant. Confocal microscopy was performed to observe the subcellular localization of ZmLDAPs and RBSDV viroplasm proteins P5-1 and P9-1 in maize protoplasts. Our results showed that P5-1 and P9-1 were distributed in the cytoplasm when expressed alone (Supplemental Fig. S5). However, when they were co-expressed with ZmLDAPs, P5-1 and P9-1 colocalized to aggregated LDs formed by ZmLDAP2, but not ZmLDAP1 (Supplemental Fig. S6).

To investigate the mechanism by which RBSDV infection utilizes LDs and associated proteins, the yeast two-hybrid (Y2H) assay was used to identify RBSDV proteins interacting with the LD proteins. The results showed that P8 interacted with ZmLDAP2, and the interaction region of P8 was the C-terminal region of 471 amino acids containing a predicted P-loop NTPase domain analyzed by Motif-scan (http://myhits.isb-sib.ch/cgi-bin/motif_scan) (Fig. 4B). To further verify this interaction, we performed glutathione-S-transferase (GST)-pull-down and luciferase complementation imaging (LCI) assays, showing that P8 interacted with ZmLDAP1 and ZmLDAP2, both *in vivo* and *in vitro* (Fig. 4C, D). When P8 was expressed alone in maize protoplasts, it localized to cytoplasm, nucleus and nucleolus (Supplemental Fig. S5), as recently reported by Li et al. (2020). Similar to RBSDV P5-1 and P9-1, P8 also co-localized with ZmLDAP2, but not with ZmLDAP1 (Fig. 4E). These results suggest that ZmLDAP2 should be a crucial factor in promoting RBSDV interaction with host LDs and a component in the viroplasm.

### Knockdown of *ZmLDAP2* suppresses RBSDV accumulation in maize

To further explore the role of ZmLDAP2 in RBSDV infection, we knocked down expression of *ZmLDAP2* in maize using the cucumber mosaic virus (CMV)-based virus induced gene silencing (VIGS) approach (Wang et al., 2016). Maize seedlings pre-inoculated with CMV-ZmLDAP2 (to silence *ZmLDAP2*) or CMV-GFP (as a control) were then inoculated with RBSDV at the two-leaf stage. Knockdown of *ZmLDAP2* expression was confirmed by RT-qPCR (Fig. 5A), and these plants did not show any developmental defects (Fig. 5B). To study whether the reduction of *ZmLDAP2* expression could regulate the synthesis of C18 PUFAs, the relative expression levels of cytoplasmic *ZmFAD*s were also detected by RT-qPCR. In CMV-ZmLDAP2 plants, there were *ca.* 70% decrease in transcription levels of *ZmFAD2a*, *ZmFAD2b* and *ZmFAD3a* (Fig. 5C). Besides, the transcript accumulation of *ZmLDAP1* was also reduced to *ca.* 50% (Supplemental Fig. S7). This result demonstrates that silencing of *ZmLDAP2* blocks LA and ALA synthesis pathways in cytoplasm.

**Figure 5.**
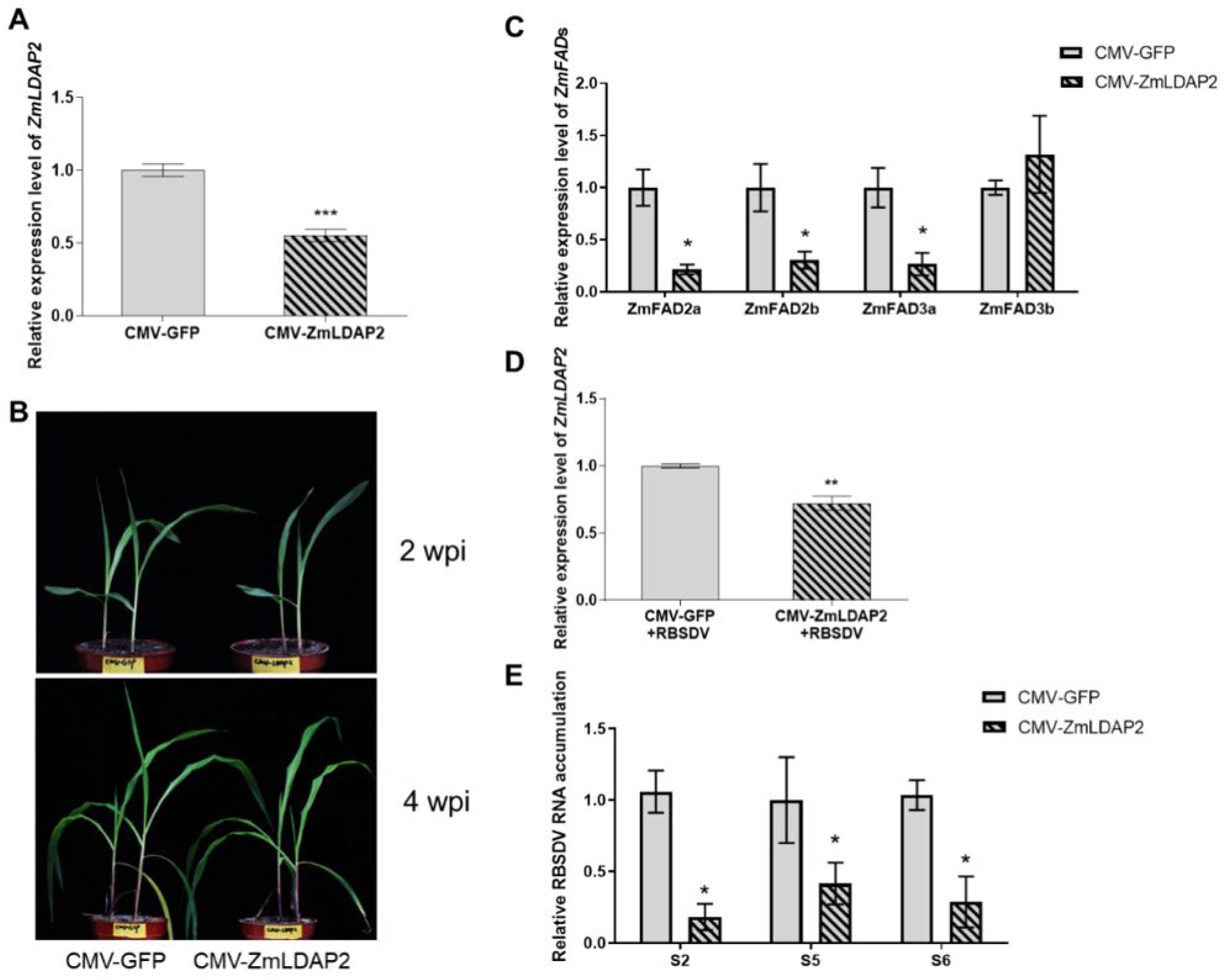
Knockdown of *ZmLDAP2* expression by CMV-VIGS impairs the cytoplasmic C18 fatty acid desaturation pathways and decreases rice black streaked dwarf virus (RBSDV) accumulation. **A**, Knockdown efficiency of *ZmLDAP2* in the third true leaves at 2 weeks post inoculation (wpi). Three independent experiments were conducted. Data were analyzed using two-tailed Student’s *t*-test and bars represent SEM (n≥4). The asterisks represent statistical significance (***, *P* < 0.001). **B,** Knockdown of *ZmLDAP2* did not cause obvious phenotype change within 4 wpi. **C,** Relative expression levels of *ZmFAD2a*, *ZmFAD2b*, *ZmFAD3a*, *ZmFAD3b* in *ZmLDAP2*-silenced maize plants and control plants. Experiments were repeated three times. Error bars represent SEM calculated from at least three maize plants (*, *P* < 0.05, **, *P* < 0.01, two-tailed Student’s *t*-test). **D,** Knockdown efficiency of *ZmLDAP2* expression at 15 dpi with RBSDV in the third true leaves. All experiments were repeated three times. Error bars represent SEM calculated from at least three replicates (**, *P* < 0.01, two-tailed Student’s *t*-test). **E,** RT-qPCR results showing the relative expression levels of RBSDV (S2, S5, and S6) in RBSDV-infected ZmLDAP2-silenced plants compared with RBSDV-infected control plants at 15 dpi. Experiments were repeated three times. Error bars represent SEM calculated from at least three plants (*, *P* < 0.05, two-tailed Student’s *t*-test).

In RBSDV-infected plants, there was *ca.* 40% decrease in *ZmLDAP2* mRNA levels in the third true leaves of the plants pre-infected with CMV-ZmLDAP2 at 15 dpi (with RBSDV) (Fig. 5D). Plants pre-infected with CMV-ZmLDAP2 supported significantly less accumulation of RBSDV RNA (*ca.* 70% reduction, Fig. 5E), in comparison with the control plants. Therefore, ZmLDAP2 plays a positive role in supporting efficient RBSDV multiplication in maize.

### RBSDV P8 and ZmLDAP2 interact with ZmPUX10

To further understand the biological functions of ZmLDAP2 and P8, these two proteins fused with GFP tag were expressed in maize protoplasts, and immunoprecipitation-mass spectrometry (IP-MS) was performed on GFP beads. ZmPUX10 was among the proteins co-precipitated with both ZmLDAP2-GFP and P8-GFP (Supplemental Fig. S8).

To validate the interaction of ZmPUX10 with ZmLDAP2, we conducted the co-immunoprecipitation (Co-IP) and LCI assays in *N. benthamiana* plants (Fig. 6A, B). Transiently expressed ZmPUX10 was found localized to LD surface and colocalized with ZmLDAP2 on LDs in maize protoplasts (Fig. 6C).

**Figure 6.**
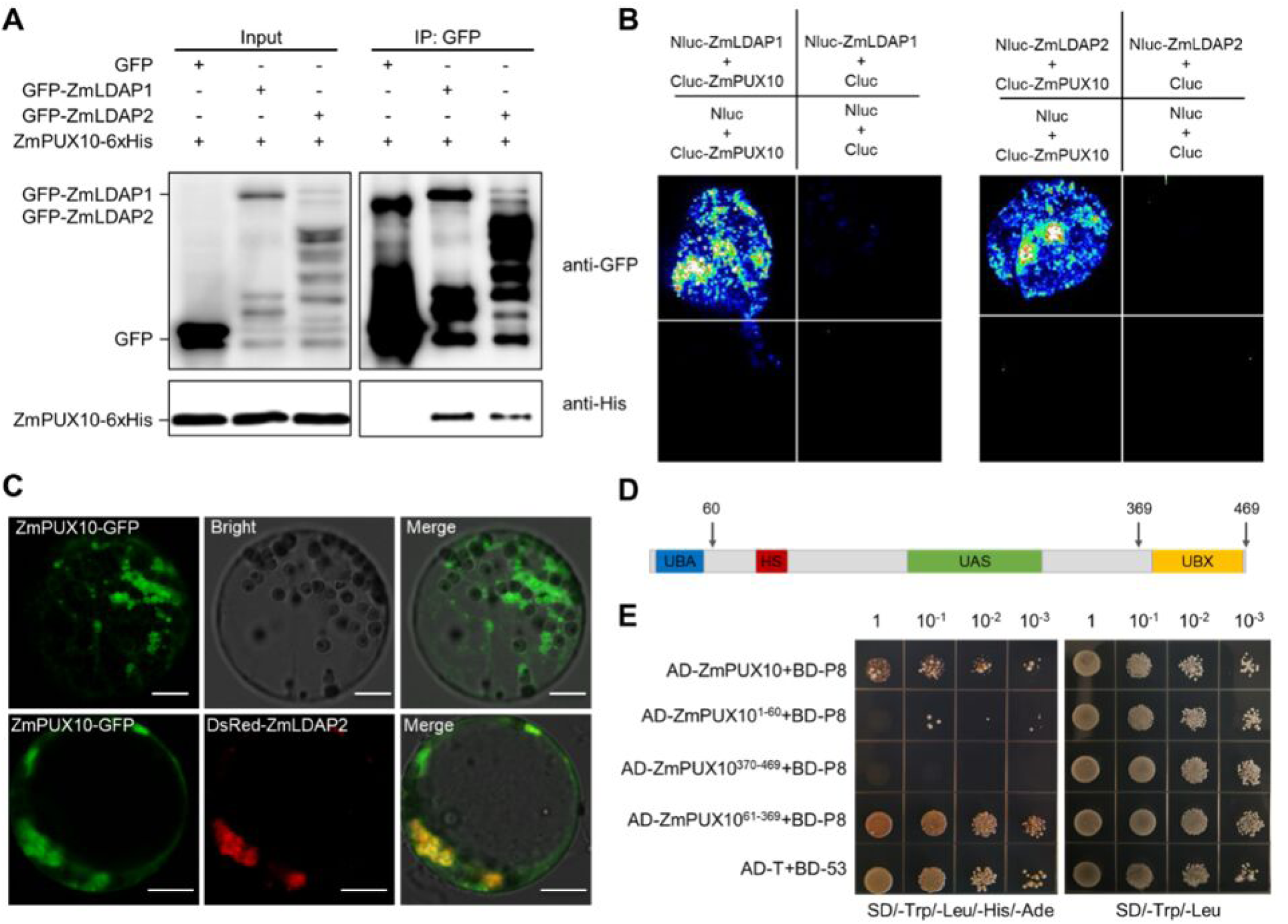
Both ZmLDAP2 and rice black streaked dwarf virus (RBSDV) P8 interact with ZmPUX10. **A,** Co-IP assays confirm that ZmPUX10 interacts with both ZmLDAP1 and ZmLDAP2 in *N. benthamiana* leaves. Total proteins were extracted and immunoprecipitated by anti-GFP beads. The coimmunoprecipitated proteins were detected by anti-His antibodies. **B,** Interactions between ZmPUX10 and ZmLDAP1 or ZmLDAP2 were detected by LCI assays in *N. benthamiana*. The *Agrobacterium* strain carrying the indicated constructs was infiltrated into *N. benthamiana* leaves which were subjected to fluorescence imaging at 3 dpai. **C,** The subcellular localization of ZmPUX10, shown by expressing GFP-ZmPUX10 alone or by co-expressing DsRed-ZmPUX10 with GFP-ZmLDAP2. Scale bars, 10 μm. **D,** The overall domain structure of ZmPUX10. The protein contains a UBA-like domain in blue (D6-P49), a hydrophobic stretch (HS) domain in red (L98-V124), a UAS domain in green (E160-Q283), and a UBX domain in yellow (P385-I467). **E,** The Y2H assay shows the interactions between P8 and ZmPUX10 truncated mutants. The combinations of constructs transformed into yeast cells were grown on selective media SD/-Trp/-Leu, and interactions were tested with SD/-Trp/-Leu/-His/-Ade. Pictures were taken after incubation at 30°C for 3 days.

Motif analysis (http://smart.embl-heidelberg.de/ and https://predictprotein.org/) revealed that ZmPUX10 also contains UBA, UBS, HS, and UAS domains similar to the published PUX10 sequences from tobacco and *Arabidopsis* (Deruyffelaere et al., 2018; Kretzschmar et al., 2018) (Fig. 6D). The Y2H assay was used to confirm the interaction between P8 and ZmPUX10, and determine the domain in ZmPUX10 responsible for their interaction (Fig. 6E). The result showed that the truncated protein without the UBA and UBX domains is sufficient to interact with P8. Obviously, these results demonstrate that ZmPUX10 interacts with both ZmLDAP2 and RBSDV P8.

### RBSDV P8 protects ZmLDAP2 from ZmPUX10-mediated degradation

Previous studies demonstrated that PUX10 channels the ubiquitinated LD proteins into the proteasome proteolytic pathway (Deruyffelaere et al., 2018; Kretzschmar et al., 2018). When the ZmLDAP1- and ZmLDAP2-expressing *N. benthamiana* leaves were treated with MG132, a specific inhibitor of the 26S proteasome, ZmLDAP2 (but not ZmLDAP1) accumulation was increased (Fig. 7A). Moreover, transient expression of ZmLDAP1 and ZmLDAP2 with ZmPUX10 in *N. benthamiana* plants at different concentrations showed that ZmLDAP2 was decreased with increasing expression of ZmPUX10 (Fig. 7B, C). Consistent with the result in Fig. 7A, ZmLDAP1 expression was not affected by ZmPUX10. Taken together, ZmLDAP2 can be degraded by 26S proteasome, and ZmPUX10 is involved in ZmLDAP2 degradation.

**Figure 7.**
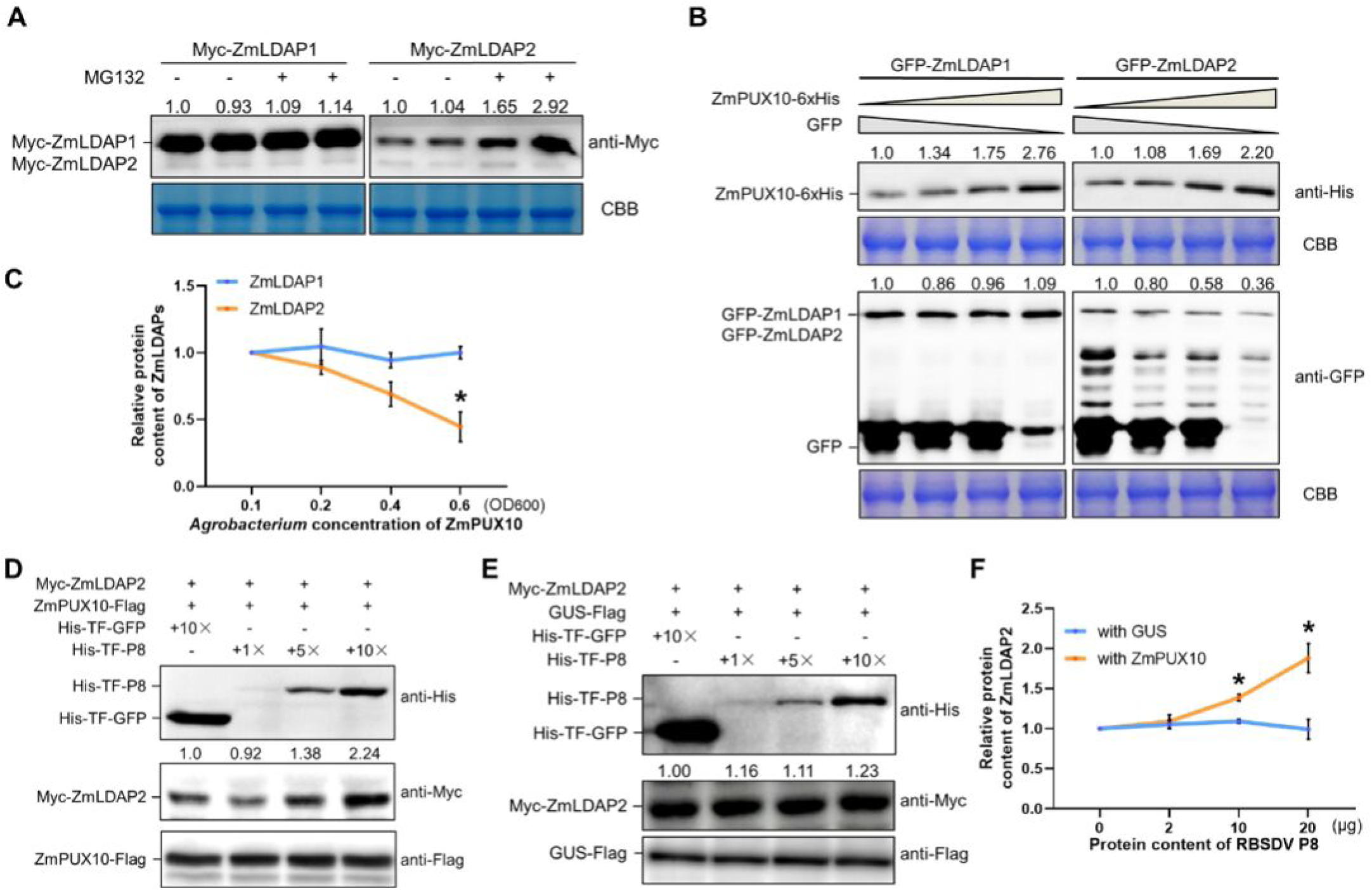
P8 protects ZmLDAP2 from ZmPUX10-mediated degradation. **A,** Degradation of ZmLDAP2 is 26S proteasome-dependent in *N. benthamiana* leaves. Myc-ZmLDAP2 or Myc-ZmLDAP1 was expressed alone in *N. benthamiana* leaves for three days. 50 μM MG132 or equal volume of DMSO (as the negative control) was infiltrated into the leaves 12 h before samples were collected. Coomassie brilliant blue (CBB) staining of rubisco was used as a loading control. The number on the panel indicates the protein level of Myc-ZmLDAP1 or Myc-ZmLDAP2 relative to the rubisco level, and the protein level with DMSO treatment in the first lane normalized to rubisco as standard 1. The experiment was repeated three times. **B,** ZmPUX10 promotes ZmLDAP2 degradation in *N. benthamiana* leaves. GFP-ZmLDAP1 or GFP-ZmLDAP2 with increasing *Agrobacterium* concentrations of ZmPUX10-6 × His (OD_600_=0.1, 0.2, 0.4 or 0.6) and decreasing *Agrobacterium* concentrations of GFP (in order to maintain the same final concentration of *Agrobacterium* in different samples) was co-infiltrated into *N. benthamiana* leaves. Infiltrated leaf areas were harvested for protein extraction at 3 dpai. The immunoprecipitated proteins were detected by either anti-His or anti-GFP antibodies. CBB staining of rubisco was used as a loading control. The number on the panel indicates the protein level of ZmPUX10-6 × His or GFP-ZmLDAPs relative to the rubisco level, and the protein level in the first lane is normalized to rubisco as standard 1. The experiment was repeated three times. **C,** Statistical analysis of the relative amount of GFP-ZmLDAP1 or GFP-ZmLDAP2 in (B). Data are shown as mean ± SEM (n = 3). (*, *P* < 0.05, two-tailed Student’s *t*-test). **D and E,** ZmPUX10-mediated ZmLDAP2 degradation is interfered by P8 in a dose-dependent manner *in vitro*. Myc-ZmLDAP2 and ZmPUX10-Flag (D) or GUS-Flag (E) were co-expressed in *N. benthamiana* leaves for three days. Equal amounts of total protein extract of *N. benthamiana* leaves were incubated with increasing amounts of His-TF-P8 (0, 2, 10, or 20 μg) or His-TF-GFP (negative control) *in vitro*. The proteins were detected by anti-His, anti-Myc or anti-Flag antibodies. The number on the panel indicates the protein level of Myc-ZmLDAP2 relative to the ZmPUX10-Flag (D) or GUS-Flag (E) level, and the protein level of the negative control in the first lane is normalized to rubisco as standard 1. The experiment was repeated three times. **F,** Statistical analysis of the relative amount of Myc-ZmLDAP2 in (D and E). Data are shown as mean ± SEM (n = 3). (*, *P* < 0.05, two-tailed Student’s *t*-test).

Moreover, to investigate whether P8 could regulate the function of ZmPUX10 on ZmLDAP2 degradation, we transiently co-expressed ZmLDAP2 and ZmPUX10 in *N. benthamiana* leaves. The same amount of total protein extracts was mixed with different amounts of purified His-TF-P8 or His-TF-GFP (negative control) and incubated for two hours, followed by Western blot analysis. The protein level of ZmLDAP2 increased with increasing amounts of His-TF-P8 (Fig. 7D, F). However, when ZmLDAP2 was co-expressed with GUS, the protein level of ZmLDAP2 was nearly unaffected by His-TF-P8 (Fig. 7E, F). These results demonstrate that P8 specially inhibits ZmLDAP2 degradation mediated by ZmPUX10.

## Discussion

The roles of the PUFAs in plant defense are complex and may depend on the particular pathogen (Choi et al., 2006; Fernández-Calvino et al., 2014; Yaeno et al., 2004). Here, we showed that RBSDV infection induces high levels of C18 PUFAs (LA and ALA) and increases transcript accumulation of cytoplasmic *FAD*s in maize leaves (Fig. 2A, B; Supplemental Fig. S3). In addition, exogenous treatment of ALA on RBSDV-inoculated maize plants resulted in aggravated dwarfing and increased viral RNA accumulation (Fig. 2C, D). As ALA is a precursor molecule in biosynthesis pathway of JA defense hormone, it is possible that the enhancement effect of ALA to RBSDV infection was induced by the JA signaling pathway (Borrego and Kolomiets, 2016). However, JA was reported to induce plant resistance to RBSDV in rice (He et al., 2017), which had the opposite effect to ALA on viral infection in this research. Hence, our results demonstrate that ALA directly promotes RBSDV infection which leads to the development of dwarfing phenotype in maize.

Overexpression of *LDAP* genes in *Arabidopsis* significantly increases LA and ALA accumulation (Gidda et al., 2016). In addition, the treatment of LA induced the proliferation of LDs in both animal and plant cells (Gidda et al., 2016; Horn et al., 2013; Jarc et al., 2018). Our results showed that silencing of *ZmLDAP2* down-regulated the transcription level of cytoplasmic *FAD*s in maize (Fig. 5C). Therefore, it is reasonable to speculate that the increased ZmLDAP2 level induced by RBSDV is able to regulate the metabolism of C18 PUFAs in the cytoplasm, and in turn, the accumulation of PUFAs in maize cells in response to RBSDV infection might enhance the biogenesis of LDs. Thus, RBSDV mobilizes cellular resources for viral multiplication both at metabolic and protein levels. However, the mechanism of ZmLDAP2-regulated *FAD* genes expression remains unknown.

Clustering of LDs is a widespread phenomenon that has been observed in previous studies. Several LD proteins of mammalian adipocytes have been shown to alter the LD morphology for their respective physiological functions. For instance, perilipin A induces the formation of LD clusters to keep the LDs from lipolysis by cytosolic lipases, which is reversed by the phosphorylation of serine 492 of perilipin A upon lipolytic stimulation (Marcinkiewicz et al., 2006). FSP27 facilitates LD clustering and then promotes their fusion to form enlarged droplets and a subsequent triglyceride accumulation (Jambunathan et al., 2011). AUP1 promotes LD clustering relying on the ubiquitination at lysine residues (Lohmann et al., 2013). Besides, HCV core protein displaces the LD protein, ADRP, from the LD surface, that results in a microtubule- and dynein-driven LD clustering towards the microtubule-organizing centers, in the perinuclear area and close to HCV replication sites (Boulant et al., 2008). In plant cells, AtLDAP1 was shown to induce LD cluster formation close to ER membranes in *Arabidopsis*, thus probably impair the dynamic intracellular movement of LDs and exchanging lipids, metabolites or proteins with other organelles (Brocard et al., 2017; Olzmann and Carvalho, 2019). Here, we characterized two LD-associated proteins of maize, ZmLDAP1 and ZmLDAP2, that showed differential transcription profiles during RBSDV infection, and induced the formation of LD clusters in maize cells (Figs. 3C, D, 4A; Supplemental Fig. S4). We demonstrate that P8 localizes to ZmLDAP2-induced LD clusters by physical interaction, as well as protects ZmLDAP2 from ZmPUX10-mediated degradation (Figs. 4B-E, 7D-F). Besides, the viroplasm proteins, P5-1 and P9-1, also targeted to ZmLDAP2-induced LD clusters in maize protoplasts (Supplemental Fig. S6), suggesting the possibility of ZmLDAP2-associated LDs needed by RBSDV replication sites. Furthermore, downregulation of *ZmLDAP2* in maize suppressed RBSDV infection (Fig. 5E). Together, these results implicate that RBSDV utilizes plant LDs (might in aggregated forms) for virus propagation, indicating the conserved functions of LDs in both animal and plant cells during viral infection. This similarity is probably due to the capability of RBSDV proliferating in both insect and plant cells. Therefore, it is necessary to further investigate the association of RBSDV with LDs in insect cells. Moreover, whether there is a similar function of LDs for other plant viruses, especially for insect-borne plant viruses, needs to be further studied, and more research is needed to advance our knowledge on the functions of LD clusters in plant cells.

LDAPs have been reported to play different roles in plant tissues. *Arabidopsis LDAP*s, *AtLDAP1*, *AtLDAP2* and *AtLDAP3*, which are the homologs of *ZmLDAP*s, showed different expression profiles in plant tissues and in response to abiotic stress (Kim et al., 2016). Here, we found that although ZmLDAP1 could also induce the formation of LD clusters, it is not targeted by any of the RBSDV proteins studied here (Fig. 4A, E; Supplemental Figs. S4, S6). In addition, although ZmLDAP1 is able to physically interact with ZmPUX10, it could not be degraded via the 26S proteasome mediated by ZmPUX10 (Figs. 6A, B, 7B). Consistent with the differential expression profiles of *ZmLDAP1* and *ZmLDAP2* in various maize tissues and in response to RBSDV infection (Fig. 3), there might be a functional specificity between these two LDAPs in both healthy and virus-infected plants. Besides, here we demonstrated that RBSDV P8 interacts with both ZmLDAP2 and ZmPUX10, and prevents ZmLDAP2 degradation via the ZmPUX10-associated pathway (Figs. 4B-D, 6E, 7D-F; Supplemental Fig. S8A), while the protein mechanism remains unclear. Some effort was made for the mechanism study, for example, a dose-dependent competitive Co-IP assay was performed to assess whether P8 interferes with the ZmLDAP2-ZmPUX10 interaction (Supplemental Fig. S9). However, the result showed that P8 did not affect the interaction between ZmLDAP2 and ZmPUX10, indicating that P8 protected ZmLDAP2 not via directly disrupting the interaction between ZmLDAP2 and ZmPUX10. Hence, the mechanism of P8 preventing ZmLDAP2 degradation needs to be further studied. Given the above, results described here help expand our understanding of physiological functions of LD proteins in plant cells during viral infection.

We have proposed a working model on the function of a lipid droplet-associated protein in RBSDV infection of maize (Fig. 8). During infection, RBSDV P8 upregulates the accumulation of ZmLDAP2 by suppressing the ZmPUX10-mediated degradation via ubiquitin-proteasome system, that results in the disturbance of the host lipid metabolism, especially C18 FA desaturation pathway. The elevated C18 PUFA level renders maize more susceptible to RBSDV infection. Therefore, our findings demonstrate that RBSDV utilizes plant LDs to facilitate infection at both protein and metabolic levels, and further studies are required to gain detailed understanding of these molecular mechanisms.

**Figure 8.**
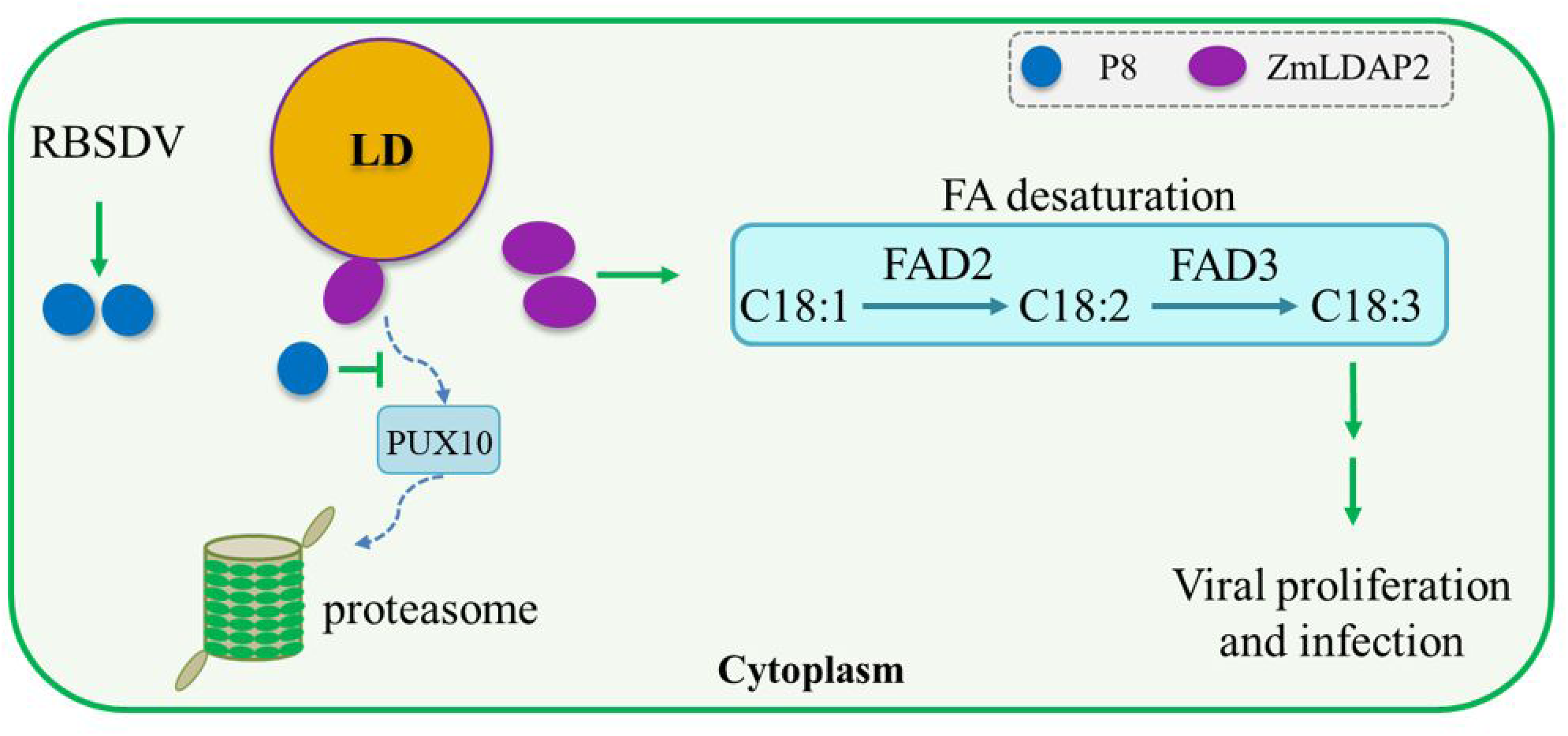
A proposed model for the function of ZmLDAP2 in RBSDV infection of maize. During RBSDV infection, P8 targets to ZmLDAP2 and protects it from ZmPUX10-mediated degradation via the ubiquitin-proteasome system, thus promoting C18 fatty acids (FAs) desaturation, which causes the accumulation of α-linolenic acid (ALA, C18:3), that facilitates viral proliferation and infection. FAD, fatty acid desaturase.

## Materials and Methods

### Plant growth and virus inoculation

*N. benthamiana* and maize (inbred line B73) plants were grown in a greenhouse (24°C, 16 h day and 22°C, 8 h night). Virus inoculation method was the same as previously described (Li et al., 2020). Maize seedlings at the two-leaf stage were fed on by viruliferous (RBSDV-carrying) or non-viruliferous (mock) small brown planthoppers (SBPH) at a ratio of 1:8 (eight insects per plant) for three days.

### Molecular cloning and plasmid construction

The coding sequences of ZmLDAP1 (accession number: NM_001320475.1, chromosome 7) and ZmPUX10 (accession number: NM_001143657.1, chromosome 1) were amplified by reverse transcription-polymerase chain reaction (RT-PCR) from maize (inbred line B73). The coding sequence of ZmLDAP2 was amplified by RT-PCR from maize (inbred line B73) is 24 nt (from nucleotides 144 to 167) short of that deposited in the GenBank (accession number: NM_001156362.1, chromosome 8). The coding sequence of RBSDV P8 (accession number: KC134296.1), P5-1 (accession number: KC134293.1), P9-1 (accession number: KC134297.1) were amplified by RT-PCR from RBSDV-infected wheat plants from Hebei Province, China. The primers used for plasmid construction in this study are listed in Supplemental Table S4.

### Sample preparation for metabolomics and GC/MS analysis

Six biological replicates of RBSDV-infected and mock-inoculated maize leaves were collected at 10 and 28 dpi, respectively, and immediately frozen in liquid nitrogen for metabolomic analysis. Metabolomic analysis was carried out following the method previously reported with slight modifications (Guo et al., 2017). In brief, 50 mg of each sample was placed in a centrifuge tube (2 mL). The samples were mixed with 10 μL adonitol (0.5 mg/mL) and 0.45 mL 75% methanol by vortexing and grinding for 5 min. After ultrasonication for 5 min and centrifugation for 15 min at 10,000 rpm, the supernatant was transferred to a fresh tube and 25 μL was taken as quality control (QC) sample. 80 μL of methoxamine hydrochloride (20 mg/mL in pyridine) was added and then incubated at 80°C for 0.5 h. Add 100 μL of the BSTFA reagent (1% trimethylchlorosilane, v/v) to the sample aliquots, incubated for 1.5 h at 70°C. Add 5 μL of fatty acid methyl esters (in chloroform) to each QC sample when cooling to the room temperature. All the samples were analyzed by GC-TOF-MS. The compounds were analyzed at Biotree Bio-technology company (Shanghai, China).

### Metabolomics data processing and statistical analysis

ChromaTOF 4.3X software (Leco Corporation) was used for raw peaks extraction, the data filtering and calibration of the baseline, peak alignment, deconvolution analysis, peak identification and integration of the peak area, and LECO-Fiehn Rtx5 database was used for metabolite identification by matching the mass spectrum and retention index. The peaks detected in less than half of the QC samples or RSD > 30% in QC samples were removed (Dunn et al., 2011). In the following, the missing value was removed and replaced with a small value that was half of the minimum positive value in the original data. Then, the data were filtered by interquantile range, and the total mass of the signal integration area was normalized for each sample. SIMCA software (V14.1, MKS Data Analytics Solutions, Umea, Sweden) was employed to run the PCA and OPLS-DA. Differentially expressed metabolites were found using Student’s *t*-test (*P* < 0.05) and OPLS-DA (variable importance in the projection, VIP > 1). Furthermore, metabolite pathways were analyzed by using the online software MetaboAnalyst (www.metaboanalyst.ca/) and KEGG (www.genome.jp/kegg/).

### RNA extraction and RT-qPCR analysis

Total RNA was extracted from maize tissue using the TRIzol reagent (Invitrogen) and subsequently treated with RNase-free DNase I (Takara Bio, Dalian, China) as instructed by the manufacturers. First-strand cDNA was synthesized with 2.0 µg RNA using M-MLV reverse transcriptase (Promega, Madison, WI, USA). RT-qPCR was performed as reported previously (Cao et al., 2012). The transcript accumulation level of *ZmUbi* (ubiquitin) was used as an internal control. The experiments were replicated at least three times. The primers used for RT-qPCR analysis are listed in Supplemental Table S4.

### ALA treatment assay

ALA (Sigma-Aldrich) was dissolved in 100% DMSO and then diluted with sterile distilled water containing 0.1% Triton X-100. DMSO was used as the control. Maize seedlings were treated by gently brushing with ALA (1 mM or 10 mM in 0.1% Triton X-100) or control (equal volume of DMSO in 0.1% Triton X-100) 24 h after virus inoculation. At least 30 seedlings were used in each treatment.

### Y2H assay

Yeast transformation was performed according to the manufacturer’s instructions (Clontech, Mountain View, CA, USA). Yeast cells (strain Y2H Gold) were co-transformed with specific bait and prey constructs. All transformants were grown on a selective medium lacking tryptophan and leucine (SD/-Trp-Leu) and a high-stringency selective medium lacking tryptophan, leucine, histidine and adenine (SD/-Trp-Leu-His-Ade) for analyzing possible interaction.

### Protein purification and *in vitro* pull-down assay

The coding sequences of *ZmLDAP1* and *ZmLDAP2* were amplified and cloned into pGEX with the GST-tag. The viral gene coding for P8 was separately cloned into pColdTF with His-tag or pET with His-tag. All recombinant plasmids were transformed into *Escherichia coli* strain BL21 (DE3) (TransGen Biotech, Beijing, China). The fusion proteins were purified with glutathione Sepharose 4B (GE Healthcare) for GST-fused proteins, or Ni-NTA Agarose (QIAGEN) for His-fused proteins according to the manufacturers’ instructions. For the pull-down assay, GST-ZmLDAP1 and GST-ZmLDAP2 were suspended on glutathione agarose resin and gently rotated at 4°C for 1 h, and then spun down to discard the supernatant. His-P8 was added into GST-ZmLDAP2-bound (or GST-ZmLDAP1-bound) beads and incubated at 4°C for 3 h. Subsequently, the beads were rinsed at least five times with IP buffer (50 mM Tris-HCl, pH6.8, 300 Mm NaCl, 1.5% glycerol, 0.6% Triton X-100, 0.1% Tween). After being eluted from beads, the proteins were detected by immunoblotting with anti-His antibodies (TransGen Biotech, Beijing, China). GST-GFP was used as the control.

### Agrobacterium tumefaciens-mediated transient expression in Nicotiana benthamiana leaves

The recombinant plasmids were introduced into *Agrobacterium* cells (strain GV3101). The transformed agrobacteria were cultured, pelleted, and then resuspended in infiltration buffer (10 mM MgCl_2_, 10 mM MES, and 100 mM acetosyringone, pH5.6). After 2 to 4 h of incubation at room temperature, appropriate agrobacterial cultures were used to infiltrate the leaves of 4-week-old *N. benthamiana* plants. Tomato bushy stunt virus p19 protein, as a strong silencing suppressor, was used to enhance protein expression. For the protein degradation assay, 10 mM MgCl_2_ buffer containing 1% DMSO (control) or an equal volume of DMSO with 50 μM MG132 (Sigma-Aldrich) for inhibition of the 26S proteasome was infiltrated into leaves 12 h before samples were collected.

### Protein extraction and immunoblotting

The protein extraction and immunoblotting assay were performed as described previously (Cao et al., 2012). Monoclonal antibodies of His and GFP (Cowin Bio, Taizhou, China) were used at a dilution of 1:5000. Monoclonal antibodies of c-Myc and Flag (Sigma-Aldrich) were used at a dilution of 1:10000.

### Maize protoplasts isolation and transfection

Maize protoplasts isolation and transfection were performed as described previously (Sheen, 1991; Zhu et al., 2014). Protoplasts were isolated from leaves of maize (inbred line Zheng 58) and transfected using a polyethylene glycol (PEG)-mediated method. The transfected protoplasts were incubated in the dark at 25°C, and harvested at 12-18 hours post transfection (hpt) and then used for further analysis.

### Visualization of LDs and protein subcellular localization

LDs were stained with Nile Red (2 μg/mL) for 10 min before visualization. The pGD vector (Goodin et al., 2002) was used for expressing recombinant proteins. For subcellular localization, plant tissues expressing GFP- or DsRed-fused proteins and stained LDs were imaged with a Leica TCS SP8 confocal microscope. The GFP, DsRed, Nile Red fluorescence were captured in the green (GFP), red (DsRed) and red (Nile Red) channels, respectively. For transient expression in *N. benthamiana* leaves, subcellular localization was determined at 3 days post agroinfiltration (dpai). For the protein expression in maize protoplasts, the fluorescence signals were visualized at 18 hpt. Three independent experiments were performed.

### CMV-VIGS in maize

CMV-VIGS assay in maize was performed as previously reported (Wang et al., 2016; Zhou et al., 2018). Briefly, *Agrobacterium* (C58C1) cultures carrying pCMV201-2B_N81_-ZmLDAP2 or pCMV201-2B_N81_-GFP was co-infiltrated with pCMV101 and pCMV301 into *N. benthamiana* leaves. At 3-4 dpai, CMV virion isolation was collected from the infiltrated leaves and subsequently vascular puncture inoculated on maize seeds (inbred line B73). The inoculated seeds were kept at 25°C in the dark for 2 days, then transferred into pots with soil and grown in a growth chamber (20°C, 16 h day and 18°C, 8 h night). These plants were subjected for RBSDV inoculation at two-leaf stage.

### IP-MS Analysis

IP-MS was performed as described previously (Wang et al., 2017) with several modifications. In brief, GFP-fused proteins were transiently expressed in maize protoplasts as described above, and samples were taken at 18 hpt. Total proteins were extracted by adding IP buffer with 1 mM DTT, 1 mM PMSF and 1% protease inhibitor cocktail and then the extracts were incubated with anti-GFP magnetic agarose beads (Lablead, Beijing, China) for one hour. The beads were subsequently washed using IP buffer for five times before stripping interacting proteins from the beads by boiling in 50 µL of 2 × sample buffer (100 mM Tris-HCl, pH6.8, 20% glycerol, 4% SDS) for 10 min. Immunoprecipitated proteins were washed by urea buffer (7 M urea, 2 M thiocarbamide) and subjected to mass spectrometry analysis. Mass spectrometry analysis was performed at the Biological Mass Spectrometry Laboratory of China Agricultural University. Matching raw MS data to peptide sequences was performed using Mascot software with the annotated proteins from Uniprot Maize database.

### Co-IP assay

The plasmids were separately transformed into *Agrobacterium* strain GV3101, and were equally mixed and co-infiltrated into *N. benthamiana* leaves. Total protein of the infiltrated leaf tissues was extracted 3 dpai as described above, and then incubated with anti-GFP magnetic agarose beads at 4°C for 0.5 h. The immunoprecipitates were detected by immunoblotting with anti-His antibodies, and GFP was used as the control. For the dose-dependent Co-IP assays, equal amounts of total proteins of *N. benthamiana* leaves expressing GFP-ZmLDAP2 and ZmPUX10-Flag were incubated with increasing amounts of purified His-TF-P8 (0, 2, 10, or 20 μg) or His-TF (negative control) *in vitro* and then immunoprecipitated by anti-GFP beads. The coimmunoprecipitated proteins were detected by anti-Flag antibodies.

### LCI assay

LCI assays were performed as previously described (Chen et al., 2008). All tested combinations were agroinfiltrated into leaves of *N. benthamiana*. The leaves were sprayed with 1 mM luciferin (Invitrogen) three days later, and then the fluorescence signal was measured by a low-light cooled CCD imaging apparatus (iXon, Andor Technology).

### Supplemental Data

The following supplemental materials are available.

**Supplemental Figure S1.** Score plot of principal component analysis (PCA) and orthogonal projections to latent structures-discriminant analysis (OPLS-DA) of metabolites from RBSDV-infected and mock-inoculated maize leaves at both infection phases.

**Supplemental Figure S2.** Up- and down-regulated metabolites of maize leaves during RBSDV infection.

**Supplemental Figure S3.** RBSDV infection regulates expression levels of *FAD* genes in maize.

**Supplemental Figure S4.** The subcellular localization of LDs, ZmLDAP1 and ZmLDAP2 in maize protoplasts.

**Supplemental Figure S5.** The subcellular localization of RBSDV P9-1, P5-1 and P8 in maize protoplasts.

**Supplemental Figure S6.** The subcellular co-localization of RBSDV P9-1 and P5-1 with ZmLDAPs in maize protoplasts.

**Supplemental Figure S7.** Relative expression levels of *ZmLDAP1* in *ZmLDAP2*-silenced maize plants and control plants.

**Supplemental Figure S8.** Representative mass spectrum for oligopeptides from ZmPUX10 protein.

**Supplemental Figure S9.** Dose-dependent Co-IP assays showing that P8 does not affect the interaction between ZmPUX10 and ZmLDAP2. GFP-ZmLDAP2 and ZmPUX10-Flag were co-expressed in *N. benthamiana* leaves for three days.

**Supplemental Table S1 (separate file).** The total primary metabolites identified in RBSDV-infected and mock-inoculated maize leaves.

**Supplemental Table S2 (separate file).** The differentially expressed metabolites between RBSDV-infected and mock-inoculated maize leaves at 10 dpi.

**Supplemental Table S3 (separate file).** The differentially expressed metabolites between RBSDV-infected and mock-inoculated maize leaves at 28 dpi.

**Supplemental Table S4.** The primers used in this study.

## Acknowledgments

We are indebted to Dr S.P. Dinesh-Kumar (University of California at Davis) for his helpful suggestions for this research. We thank Dr Zhen Li in the Biological Mass Spectrometry Laboratory of China Agricultural University for providing helps in IP-MS data measure. We thank Dr Andrew Jackson (University of California, Berkeley) for providing the pGD vector, Dr Jianmin Zhou (Institute of Genetics and Developmental Biology, Chinese Academy of Sciences) for providing the plasmids for luciferase complementation imaging assay.

